# NAD^+^-dependent deacetylase SIRT1 is essential for meiotic progression and controls repair-recombination efficiency

**DOI:** 10.1101/666891

**Authors:** Harshita Kaul, Shaunak Deota, Amit Fulzele, Anne Gonzalez-de-Peredo, Ullas Kolthur-Seetharam

## Abstract

Meiotic components and their functions have been extensively studied. Yet, the interplay between molecular factors and regulation of their functions that is brought about by post-translational modifications, specifically (de)-acetylation, is not well characterized. SIRT1, a NAD^+^-dependent deacetylase has been previously shown to be necessary for spermatogenesis. However, whether it has any role to play in mammalian meiosis remains to be uncovered. Our findings identify SIRT1 as a key determinant of meiotic progression. Knocking out SIRT1 specifically in meiocytes (SIRT1*^Δmeio^*) led to a delay in progression through pachytene and repair of double strand breaks. Interestingly, despite these deficits, meiotic loss of SIRT1 did not affect synapsis nor did it lead to pachytene arrest or apoptosis. Moreover, our results demonstrate that SIRT1 is required for regulating crossover frequency and its absence results in higher crossover events. Therefore, our study brings to the fore a novel regulatory factor/mechanism that is necessary for coupling of synapsis and recombination. This is noteworthy since mutations in core meiotic components result in gross defects in synapsis, repair and recombination, and very few studies have reported the differential regulation of these processes. Further, exposing SIRT1*^Δmeio^* to low/moderate doses of ©-irradiation indicated that SIRT1 might be involved in eliciting recombination checkpoint arrest and in its absence pachytene cells progress to diplotene stage, unlike in the SIRT1*^WT^* mice. Importantly, exogenous damage resulted in enhanced retention of ©H2AX in SIRT1*^Δmeio^* diplotene cells, reiterating the critical role that SIRT1 plays in regulating repair efficiency/kinetics. Molecularly, we find that SIRT1 interacts with MRN complex and lack of SIRT1 causes hyperacetylation of several non-histone proteins including the MRN components. Given that SIRT1*^Δmeio^* mice mimic MRN hypomorphs, we propose that SIRT1-dependent deacetylation of these proteins is crucial for normal meiotic progression. Taken together, our study uncovers a previously unappreciated role of SIRT1 in meiotic progression.

**Author Summary:** Meiosis is a key process in germ cell development that is essential for generating genetic diversity via recombination. It involves precise spatio-temporal orchestration of various molecular events such as chromosomal synapsis, repair and recombination. Whereas the core meiotic components are well known, upstream factors that might be important for regulating their functions and also couple the downstream processes are less explored. In this paper, we report that SIRT1, a NAD^+^-dependent deacetylase, is necessary for meiotic progression by identifying its role in coupling of synapsis and recombination. By generating a meiosis specific knockout of SIRT1, we show that its absence in spermatocytes leads to inefficient/delayed repair and progression through pachytene. We have also uncovered that SIRT1 exerts control over recombination (cross over) frequency. Interestingly, our findings demonstrate that SIRT1 provides protection against exogenous genotoxic stress possibly by eliciting meiotic checkpoints. Thus, this study provides both cellular and molecular insights into the importance of SIRT1 mediated protein deacetylation in governing meiosis in mammals.

## Introduction

Spermatogenesis is a highly orchestrated process of germ cell development involving meiotic and post-meiotic events, which are intimately linked to genome and chromatin reorganization [1]. In addition, progression through these stages is intrinsically coupled to mechanisms that elicit checkpoints, in response to both endogenous and exogenous stresses such as DNA damage [2, 3]. Importantly, meiotic recombination entails programmed induction of double strand breaks (DSBs), whose repair and choice of resolution of the double Holliday junctions via either crossover or non-crossover events determines the recombination frequency and thus the final outcome of meiosis [3-6]. Although, chromatin and non-chromatin players that impinge on these processes are known, their regulation by post-translational modifications is poorly characterized [7-16]. Specifically, importance of de-/acetylation dependent control of meiosis has not been elucidated thus far.

Interestingly, previous reports including from our lab have demonstrated that SIRT1 (NAD^+^-dependent deacetylase) is abundantly expressed during spermatogenesis, more so in meiocytes [17, 18]. In addition, we have recently identified a shorter isoform, which lacks a domain that imparts substrate specificity and is predominantly expressed in the testis [18]. Not surprisingly therefore, various models of SIRT1 loss of function and testis-specific conditional mutants have been shown to cause sterility [17, 19-24]. While many of these reports have provided insights into the role of SIRT1 in post-meiotic maturation [17, 22], its relevance during meiosis has not been addressed. Although, knocking out *Sirt1* using *Stra8-Cre* led to abnormal spermatogenesis and reduced fecundity, any potential meiotic defects were poorly characterized [17]. Importantly, perturbing SIRT1 expression or function in testis resulted in loss of pachytene cells, indicating a plausible role for this protein in orchestrating progression through meiosis [17, 20]. This is particularly relevant since SIRT1 expression is highest in meiotic prophase [17, 20] and it has been otherwise shown to be involved in DSB repair in somatic cells [25-31]. Therefore, if/how SIRT1 affects spermatocytes at cellular and molecular levels remains unknown.

Seminal studies have identified key components of the meiotic machinery, which are essential for efficient DNA damage, repair and recombination. Mutating components such as SPO11, ATM, TRIP13, DMC1 and MLH1 leads to meiotic arrest, loss of meiocytes and therefore sterility [32-38]. Despite these, the molecular basis for functional interactions between many of these factors is less understood. For example, mice harboring hypomorphic alleles of the MRN complex are sub-fertile and have meiotic deficits without a change in meiotic population [39, 40], hinting at perturbations of certain molecular interactions/functions that result in such a phenotype. Further, recent reports employing combination mutants of molecular factors, which are essential for ensuring progression through meiotic stages, have provided interesting insights into possible regulatory loops and checkpoints exerted by them. Notably, perturbations involving MRN hypomorphs (*Mre11^ALTD/ALTD^* and *Nbs1^ΔB/ ΔB^*) [39, 40] or *Atm^−/−^*; *Spo11^+/−^*; *Trip13^mod/mod^* [41-43] indicated that fine-tuning of activities of these proteins is critical in coupling different molecular processes such as synapsis, repair and recombination. These highlight the need for further studies that will not only unravel functional interactions between core meiotic components, but also of efforts to identify upstream regulators. Importantly, it is intuitive to expect that post-translational modification/s (PTM/s) based regulation would exert control over molecular-/temporal-coupling of these processes and hence define fidelity of meiosis.

In this study, we describe the key role that SIRT1 plays in regulating meiotic progression. Our findings demonstrate that loss of SIRT1 in meiocytes affects efficiency of repair and recombination, without causing an arrest in spermatogenesis. Notably, *Sirt1^Δmeio^* mice show a delay in repair and increased cross over frequencies. Exposing these mice to ionizing radiation also revealed that SIRT1 is necessary to elicit checkpoints in response to mild genotoxic stress. Together, this report identifies SIRT1, a NAD^+^-dependent deacetylase, as a critical meiotic regulator that is required to couple molecular processes with cellular progression through meiosis.

## Results

### Meiotic loss of SIRT1 leads to hyper-acetylation of proteins

Although, previous reports have alluded to a role of SIRT1 in meiosis during spermatogenesis, the precise function of this NAD^+^-dependent deacetylase during meiotic progression is still unknown. Thus, we set out to determine the phenotype of meiocyte-specific loss of SIRT1 by crossing *SirT1 Exon-4^lox/lox^* mice with *Spo11-Cre* mice, henceforth called *Sirt1^Δmeio.^* (Figs 1A, 1B, S1A and S1B). To check for changes in acetylation of histone and non-histone proteins, we used both site specific and pan anti-acetyl antibodies. Consistent with earlier reports on the effect of loss of SIRT1 in mammals [44], we did not see global changes in histone acetylation (Figs 1C and 1D and S5 Fig), and it is most likely that levels of H3K9Ac and H4K16Ac (sites targeted by SIRT1) could be altered at specific loci. However, we found dramatic changes in acetylation of several non-histone proteins in testis (Figs 1E and 1F). These clearly show that loss of SIRT1 indeed leads to global hyper-acetylation of proteins in the germ cells.

**Figure 1.**
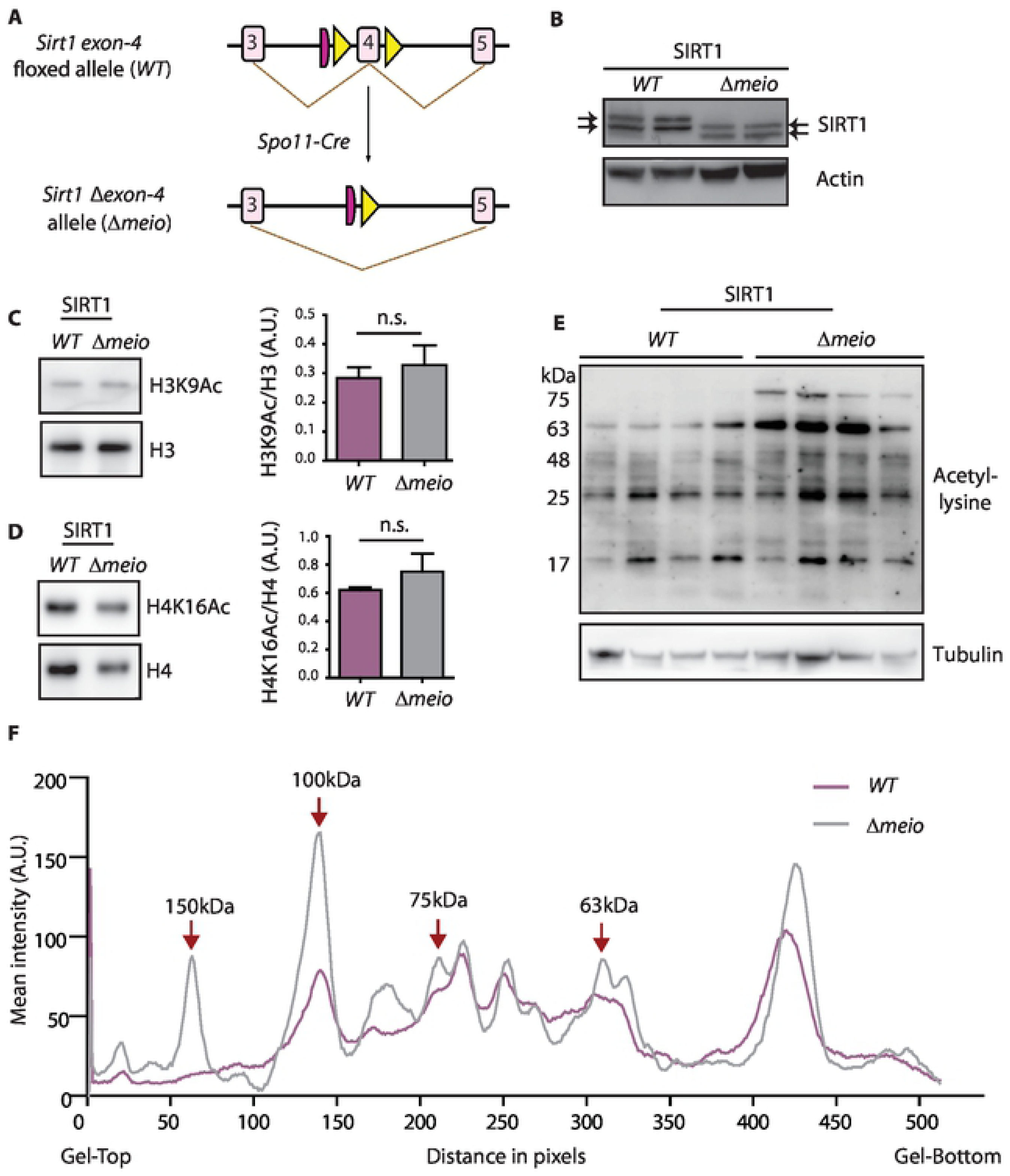
Hyper-acetylation of proteins in *Sirt1^Δmeio^*. (A) Schematic of the strategy for generating *Sirt1^Δmeio^* mice. (B) Immunoblot of SIRT1 from testis lysate of *Sirt1^WT^* and *Sirt1^Δmeio^* mice. Arrows point to two isoforms of SIRT1, which have exon-4 excision. (C, D) Representative immunoblots of (C) H3K9Ac and (D) H4K16Ac in acid-extracted histones from testis and quantifications from triplicate samples. Students t-test done to determine statistical significance (E) Immunoblot of testis lysates from *Sirt1^WT^* and *Sirt1^Δmeio^* mice using with pan anti-acetyl lysine antibody. (F) Mean intensity profile of acetylated protein bands, from an independent immunoblot, from *Sirt1^WT^* and *Sirt1^Δmeio^* mice testis lysates. Hyperacetylated bands at the mentioned molecular weights, based on mobility, are indicated. Plot showing mean of four samples per genotype.

### Meiotic populations and synapsis are unaffected in *Sirt1^Δmeio^* mice

Unlike earlier reports of loss of SIRT1 either in the whole body or during pre-meiotic stages [17, 20, 24], we did not see any alterations in different cell populations, gross abnormalities in seminiferous tubule morphology or apoptotic cells in *Sirt1^Δmeio^* mice (S1C-S1E Figs). Further, on assessing the phenotype of mice reared at two separate animal facilities (AH-1 and AH-2, described in Methods), and as illustrated in the rest of the paper, the effect of loss of SIRT1 on meiotic progression was largely indistinguishable between these cohorts (S1C Fig).

Chromosome spreads from *Sirt1^WT^* and *Sirt1^Δmeio^* testis were stained for synaptonemal complex proteins SYCP3 and SYCP1, which mark the lateral and axial elements, respectively, to assess meiotic progression and synapsis. We found similar number of cells in leptotene, zygotene, pachytene and diplotene stages between *Sirt1^WT^* and *Sirt1^Δmeio^* mice, both in young adults and during first wave of spermatogenesis (Figs 2A-2D). Importantly, unlike the phenotypes observed earlier when *Sirt1* was knocked out in pre-meiotic stages [17, 22], we did not see any change in the relative percentage of pachytene cells. Further, we also found no abnormalities in terms of either desynapsis or breaks, both in autosomes and in sex chromosomes. SMC3β staining patterns were similar between *Sirt1^WT^* and *Sirt1^Δmeio^* mice (S2A Fig), indicating no difference in sister chromatid cohesion. In addition, scoring for H3K9Me3 and TRF1 did not indicate any centromeric or telomeric fusions, respectively (S1F and S1G Figs). However, it was interesting to see that *Sirt1^Δmeio^* meiocytes exhibited a decrease in Synaptonemal Complex (SC) length, and a skew in the relative percentages of early and late pachytene cells (Figs 2E and 2F). Together, these indicated that despite having no gross defects on meiotic populations and synapsis between homologous chromosomes, loss of SIRT1 during meiosis resulted in delayed progression through pachytene.

**Figure 2.**
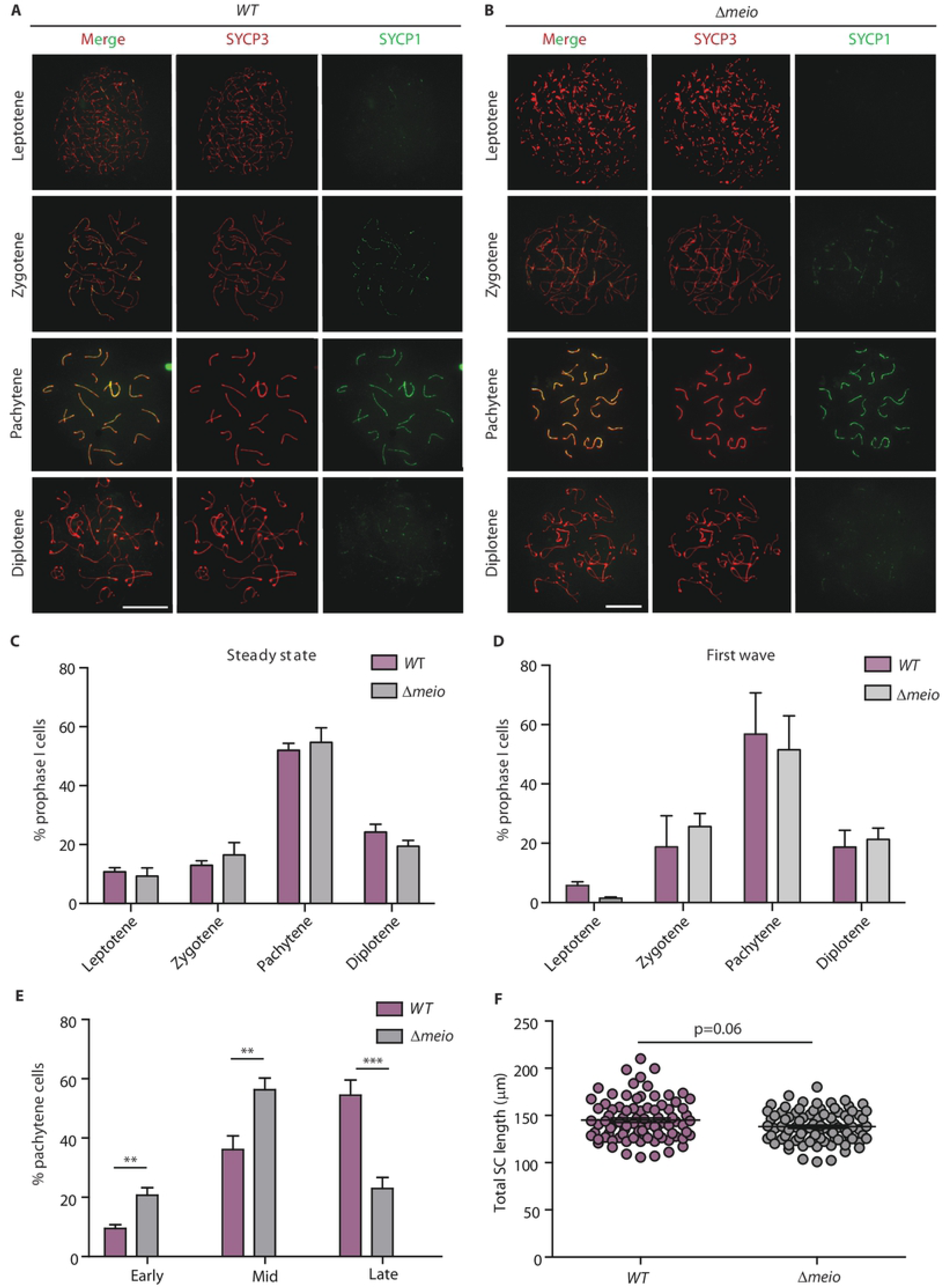
Distribution of meiotic populations and synapsis is unaffected in *Sirt1^Δmeio^* spermatocytes. (A, B) Progression of meiosis in mice at 24 d.p.p., assessed by staining with SYCP3 and SYCP1 in (A) *Sirt1^WT^* and (B) *Sirt1^Δmeio^* mice. Scale bar: 10μm. (C, D) Distribution of cells at different prophase I stages, (C) at steady state (from 5 weeks old mice) and (D) at first wave (from 24 d.p.p. mice). Mean ± SEM, N=4 mice per genotype, n=563 for *Sirt1^WT^* and n=616 for *Sirt1^Δmeio^*. (E) Distribution of pachytene cells at early, mid or late stages. (F) Average autosomal SC length at late pachytene (MLH1 positive) cells. Mean ± SEM, N=4 mice per genotype, n=86 for *Sirt1^WT^* and n=88 for *Sirt1^Δmeio^* mice. Students t-test and Mann-Whitney U test done to determine statistical significance in Fig 2E and 2F respectively.

### Absence of SIRT1 results in abnormal retention of γH2AX in pachytene stage

Although, a lack of cellular phenotype was surprising, particularly since SIRT1 is most abundant in meiotic cells, we scored for possible defects in double strand break (DSB) induction, repair and recombination efficiencies in *Sirt1^Δmeio^* mice. The process of meiosis begins with genome-wide programmed DSB formation, characterized by appearance of ©H2AX during leptotene and zygotene, and its clearance from autosomes is used as a readout for efficient repair [45]. Either abnormal levels or retention of ©H2AX, indicative of delayed or defective repair, is often associated with altered meiotic progression. In order to study the effect of loss of SIRT1 on induction or resolution of DSBs, we stained meiotic chromosome spreads for ©H2AX. As seen in Figs 3A and 3B, ©H2AX staining was similar at leptotene stage in both wild type and *Sirt1^Δmeio^* mice. Interestingly, while we saw an abnormal retention of ©H2AX in pachytene cells from *Sirt1^Δmeio^* mice as compared to *Sirt1^WT^*, it was cleared by diplotene (Figs 3A and 3B and S3A and S3B Figs).

**Figure 3.**
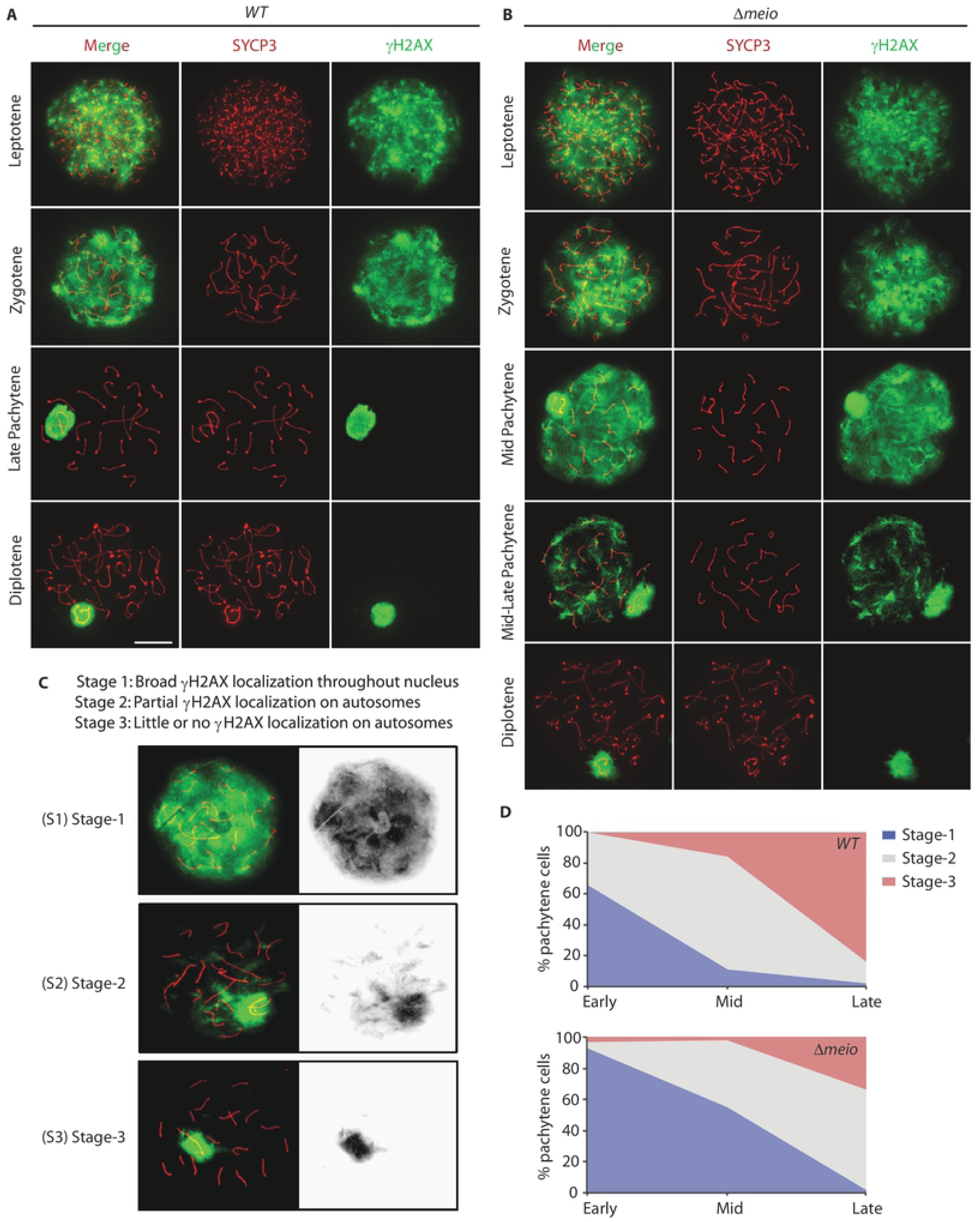
Abnormal retention of γH2AX in *Sirt1^Δmeio^* mice. (A-B) Immunostaining for SYCP3 and ©H2AX showing abnormal retention of ©H2AX patches in late pachytene stage in *Sirt1^Δmeio^* spreads (B). (C) Representation of the scheme for classification of pachytene cells based on ©H2AX pattern on autosomes, adapted from *Abe H. et. al.* [46]. (D) Integrative analyses of spermatocytes with ©H2AX staining from *Sirt1^WT^* and *Sirt1^Δmeio^* mice; as described in (C). N=4 per genotype. n=308 for *Sirt1^WT^* and n=384 for *Sirt1^Δmeio^* mice.

To get a quantitative measure, we binned pachytene cells into three distinct categories vis-à-vis ©H2AX levels/pattern, as described by a previous report [46]. Specifically, cells were scored as belonging to Stage-1, −2 or −3 based on whether ©H2AX staining was cloud like, was in patches or was completely cleared from autosomes, respectively (Fig 3C). Notably, we found that *Sirt1^Δmeio^* germ cells were represented more in Stage-1 compared to *Sirt1^WT^* cells both at early- and mid-pachytene, and in Stage-2 at late-pachytene (Fig 3D and S3C-S3E Figs). Moreover, mice housed at AH-1 and AH-2 phenocopied each other vis-à-vis abnormal retention of ©H2AX in *Sirt1^Δmeio^* mice (S3C-S3H Figs). We specifically highlight that independent of the underlying mechanism; abnormal retention of ©H2AX clearly indicated abrogated DSB homeostasis, which could have been caused by either increased DSB formation or a delay in repair in *Sirt1^Δmeio^* mice.

### Delayed repair kinetics in *Sirt1^Δmeio^* mice

In order to determine whether the abnormal ©H2AX pattern was due to an increase in the number of DSBs or delayed repair, we assayed for homologous recombination repair markers like Replication Protein A (RPA32). The localization pattern and number of RPA foci on SC axis across meiotic stages are highly regulated and are used as *bona-fide* markers to assess repair kinetics/efficiency [47, 48]. We observed that while the numbers of RPA foci on autosomes in the early pachytene stage were similar in *Sirt1^WT^* and *Sirt1^Δmeio^* cells (Figs 4A and 4B), foci counts at late pachytene were 3-fold higher in *Sirt1^Δmeio^* cells when compared to *Sirt1^WT^* (Figs 4C and 4D). Consistent with disappearance of ©H2AX, RPA foci were eventually cleared by diplotene. We also observed a similar pattern for pRPA foci in *Sirt1^Δmeio^* cells (S4 Fig). These results clearly indicate that while loss of SIRT1 did not have any effect on DSB induction, repair kinetics/efficiency was affected and importantly was decoupled with synapsis and progression through meiosis. The delay in repair may be due to delayed loading or reduced activity of downstream mediators of HR repair or a direct effect of deacetylation of ©H2AX itself. However, based on our acetylation and SIRT1-interactome results (Fig 1E and Fig 6A), it is more likely to be caused by non-histone mediators of repair.

**Figure 4.**
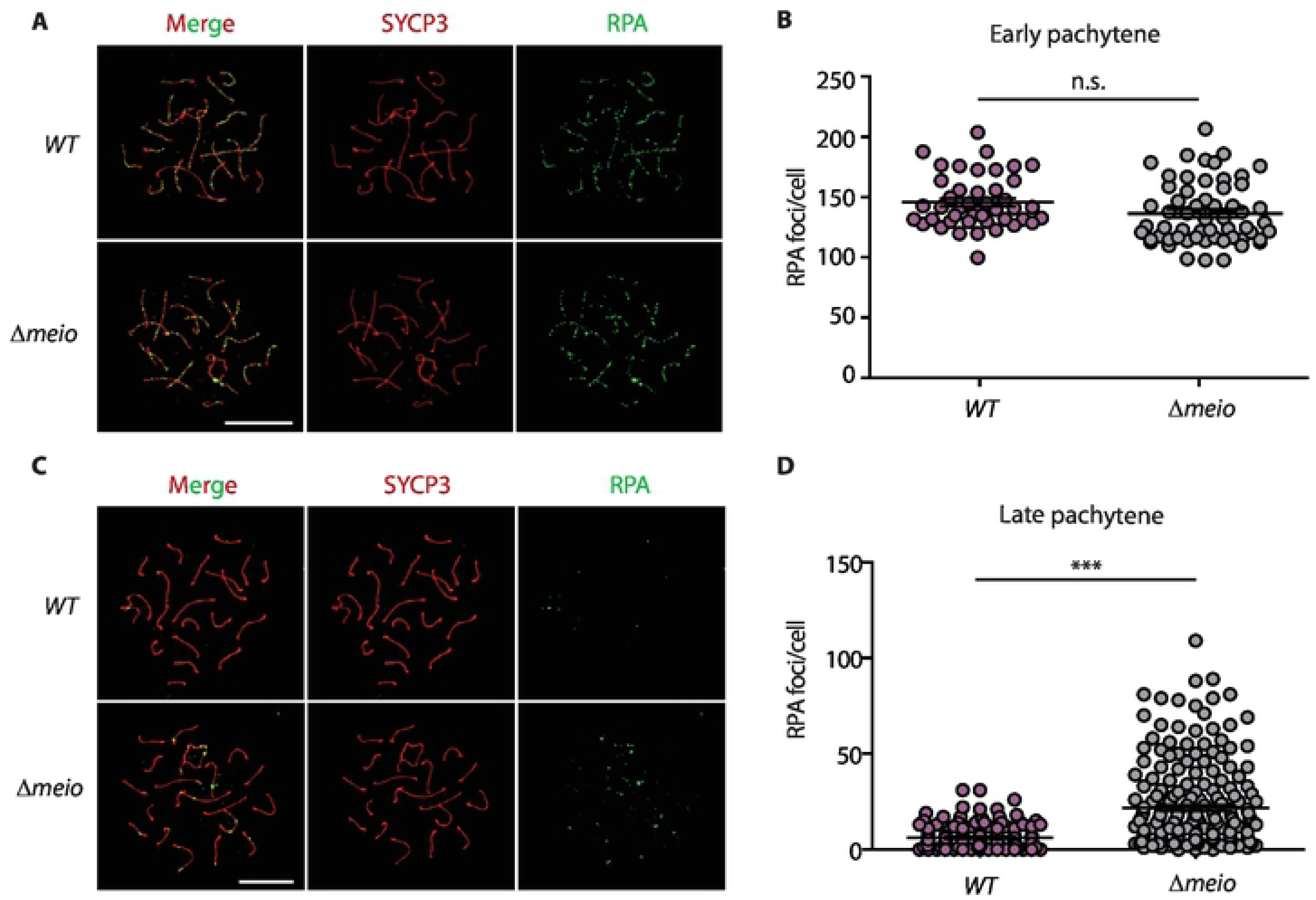
Meiotic loss of SIRT1 causes persistence of DSB repair intermediates in pachytene cells. (A, C) Representative images of cells immunostained for SYCP3 (red) and RPA32 (green) at (A) early and (C) late pachytene stages. Scale bar 10μm. (B, D) Quantification of number of RPA foci co-localizing with SYCP3 axis at (B) early and (D) late pachytene stages. Mean ± SEM, N= 5 per genotype, n=44 for *Sirt1^WT^* and n=58 for *Sirt1^Δmeio^* mice for early pachytene, and n=142 for *Sirt1^WT^* and n=204 for *Sirt1^Δmeio^* mice for late pachytene. Quantitations from mice at both AH-1 and AH-2. Students t-test used for determining statistical significance.

### SIRT1 affects crossover frequency

Given that DSB repair kinetics/efficiency has been linked to crossover frequency [49, 50], we wanted to determine if meiotic SIRT1 loss had any impact on recombination. We assessed the crossover (CO) frequency using MutL Homolog 1 (MLH1) as a marker and observed a significant increase in the average number of MLH1 foci in *Sirt1^Δmeio^* compared to *Sirt1^WT^* (Figs 5A and 5B). There was an increase in the number of bivalents with two and three MLH1 foci (Figs 5D and 5E). We also saw a high percentage of cells with one MLH1 focus on sex body in *Sirt1^Δmeio^* compared to *Sirt1^WT^* mice (Fig 5C). Additionally, we did not observe any achiasmate cells in metaphase spreads from *Sirt1^Δmeio^* mice (S2B Fig). Thus, our results indicated that SIRT1 impinges on meiotic cross over frequency, which was hitherto unknown.

**Figure 5.**
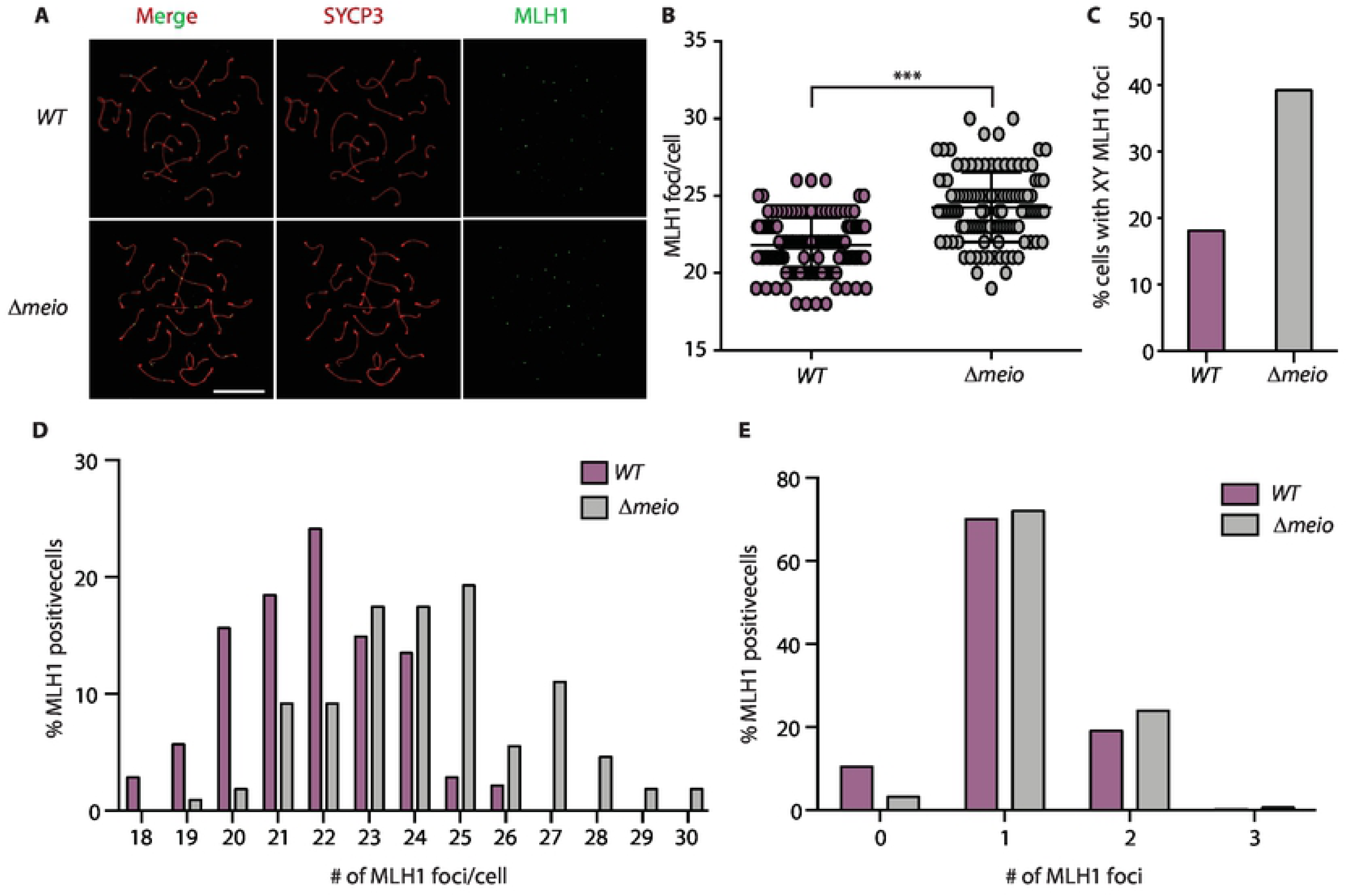
SIRT1 is required for normal cross over frequency. (A) Representative images of cells immunostained for SYCP3 (red) and MLH1 (green) at late pachytene stage. Scale bar 10μm. (B). Quantification of the total number of autosomal MLH1 foci per pachytene cell showing an increase in *Sirt1^Δmeio^* cells. Mean ± SEM, Mann-Whitney U test done to determine statistical significance. (C) Percentage of MLH1 positive XY chromosomes from *Sirt1^WT^* and *Sirt1^Δmeio^* pachytene spermatocytes. (D) Frequency distribution of MLH1 foci at late pachytene. (E) Percentage of bivalent autosomes with the indicated numbers of MLH1 foci. N= 4 per genotype, n=94 from *Sirt1^WT^* and n=102 from *Sirt1^Δmeio^* mice.

### SIRT1 is associated with the MRN complex in testis

The results presented above clearly illustrated inefficient repair/recombination in the absence of SIRT1 functions in spermatocytes. To gain preliminary mechanistic insights, we used proteomics to map SIRT1-interactome in the testis, specifically to look for factors that could explain the phenotype of *Sirt1^Δmeio^* mice. Interestingly, we found that SIRT1 was associated with the MRN complex in the testis (Figs 6A-6C). Although, SIRT1 has been implicated in regulating MRN complex via NBS1 in somatic cells [28, 51], whether it interacts with and deacetylates other components in the complex, and more so during meiosis, has not been investigated until now. We also found MRE11 to be hyperacetylated following SIRT1 inhibition in HEK293T cells (S5 Fig). Even though the data presented in S5 Fig was obtained from heterologous cell line, it nevertheless shows for the first time that MRE11 acetylation is SIRT1-dependent. Further, earlier reports have also indicated that RAD50 acetylation is increased upon SIRT1 loss of function [52, 53]. In this regard, our findings on MRE11 hyperacetylation are significant and together suggest that SIRT1 potentially regulates all the components of the MRN complex. Moreover, analysis of SIRT1 interacting proteins from testis using Gene Ontology (GO) functional analyses and STRING database showed that other key regulators of repair/recombination, which are associated with MRN complex, could be involved in a functional network (Figs 6A-6C). Even though we could immunoprecipitate MRN components from human cells, our efforts to check hyperacetylation of immunoprecipitated MRE11 and RAD50 from spermatocytes of *Sirt1^Δmeio^* mice failed, possibly due to inefficient pull-down or relative lower abundance of endogenous proteins in mice testis (Data not shown). Although speculative, the hyperacetylated bands in *Sirt1^Δmeio^* testis correspond to the molecular weights of MRE11, NBS1 and RAD50 (Fig 1F). It should also be noted that the phenotype we have described here mimics that reported in *Mre11^ALTD/ALTD^* and *Nbs1*^⊗B/ ⊗B^ hypomorphs to a large extent [39]. Thus, in the future it will be exciting to tease out the contributions of each of these SIRT1-dependent de-/acetylations in regulating interactions and/or activities of MRE11, RAD50 and NBS1 (Fig 6D) using acetyl-/deacetyl-mimic mutant versions of these proteins specifically expressed during meiosis.

**Figure 6.**
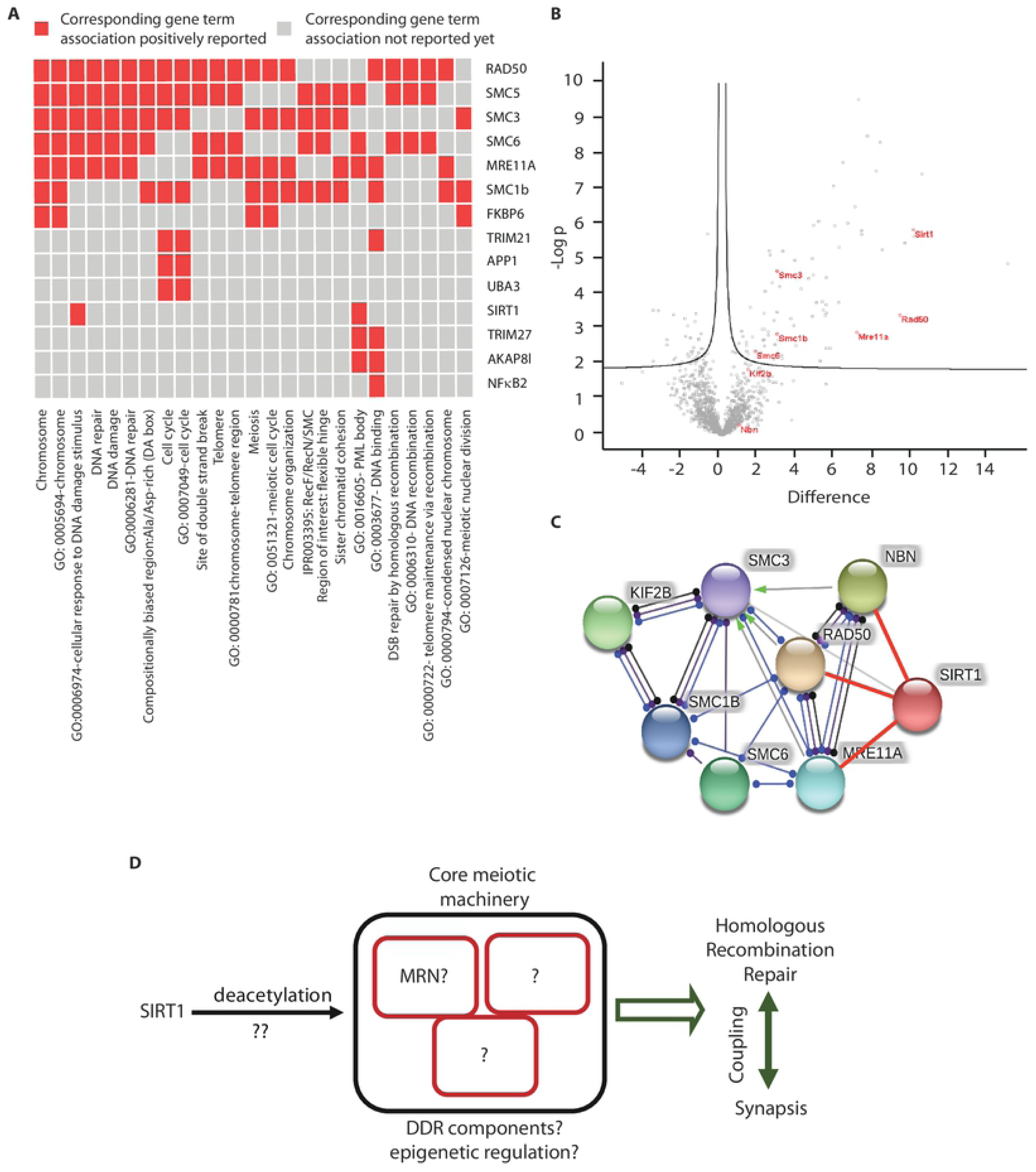
SIRT1 interactome in the testis reveals MRN components. (A) GO analysis of the proteins interacting with SIRT1 in testis. (B) Volcano plots depicting SIRT1 interacting proteins involved in meiosis (red). (C) STRING database analysis showing SIRT1 interaction network of, MRN/associated factors. (D) Proposed model describing the role of SIRT1 in regulating meiotic repair, recombination and progression possibly brought about by deacetylation of non-histone proteins that are key determinants of meiosis, as indicated.

### *Sirt1^Δmeio^* mice are hypersensitive to low exogenous DNA damage

*Sirt1^Δmeio^* mice showed delayed repair and progression through pachytene without any effects on synapsis or distributions of meiotic populations. This prompted us to investigate if an absence of SIRT1 would result in hyper-sensitization to low/moderate levels of genotoxic stress. In particular, we wanted to address if SIRT1 had any role to play in eliciting meiotic checkpoints in response to exogenous DNA damage.

Previous reports have indicated that irradiation (IR) of C57BL/6 mice with doses below 8Gy does not lead to any apparent apoptosis of germ cells or reduction in sperm counts, and spermatogenesis proceeds uninterrupted [54]. It is interesting to note that across multiple doses of IR, we saw an increased global accumulation of ©H2AX. Specifically, irradiation with a moderate dose of 3Gy showed that the percentage of cells displaying pattern characterized as stage-1 was significantly higher in *Sirt1^Δmeio^* cells (Fig 7B and S6A-S6C Figs). Moreover, the severity of the phenotype vis-à-vis retention of ©H2AX was directly correlated to the extent of exogenous damage and was significantly more than what was observed in undamaged *Sirt1^Δmeio^* mice (Fig 7B and S6A-S6F Figs).

**Figure 7.**
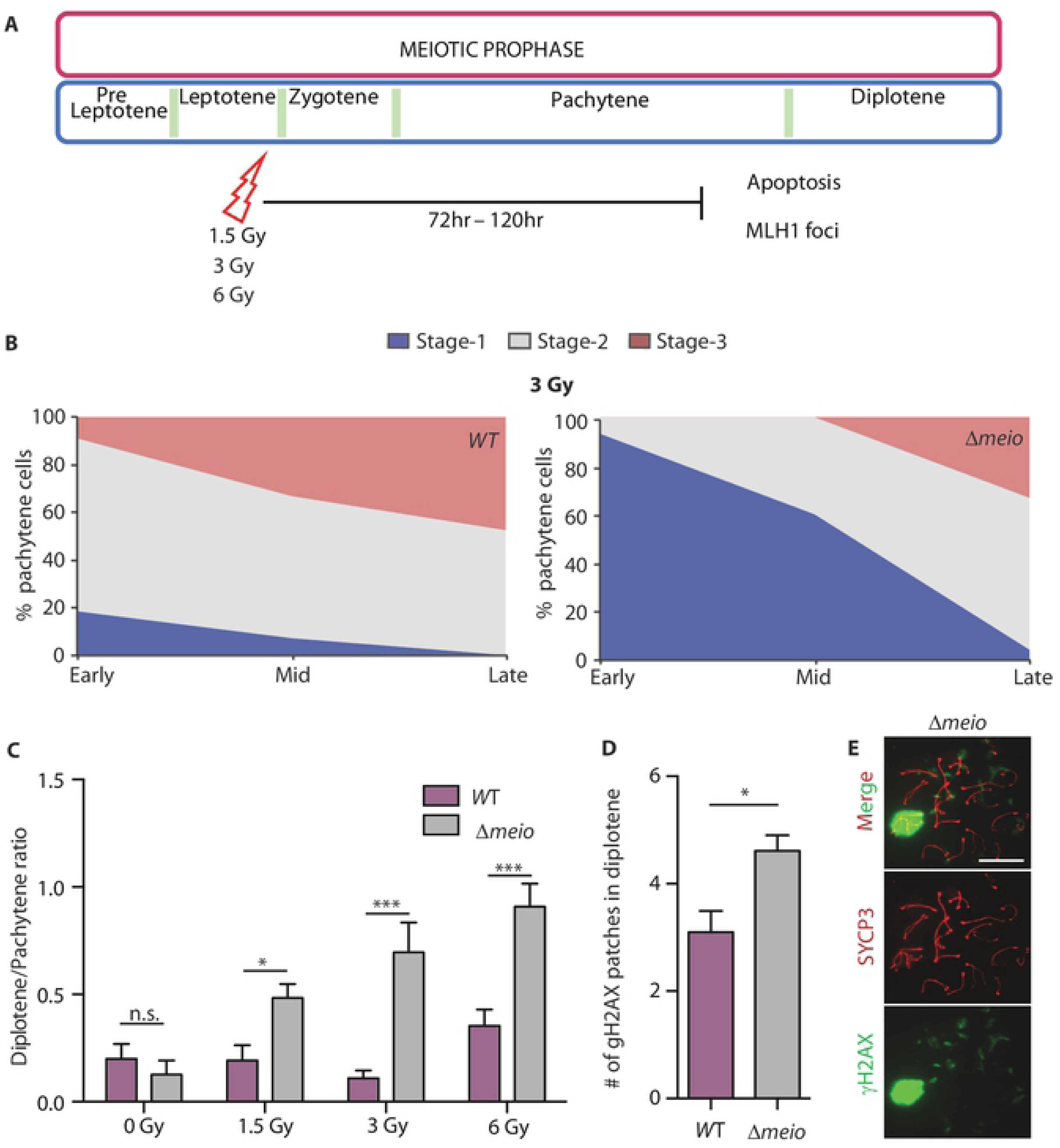
SIRT1 elicits checkpoint response and its loss results in hypersensitization of meiocytes to exogenous damage. (A) Schematic of the experimental paradigm followed for gamma irradiation of *Sirt1^WT^* and *Sirt1^Δmeio^* mice. (B) Integrative analyses ©H2AX staining in spermatocytes upon ©-irradiation, as described in Figure 2C. N=3 per genotype, n=294 for *Sirt1^WT^* and n=239 for *Sirt1^Δmeio^*. (C) Ratio of diplotene:pachytene stages following ©-irradiation, as indicated. (D) Quantification of the number of ©H2AX patches in diplotene cells following ©-irradiation. Mean ± SEM, N=3 per genotype, n=20 for *Sirt1^WT^* and n=102 for *Sirt1^Δmeio^*. (E) Representative images of cells immunostained for SYCP3 (red) and ©H2AX (green) at early diplotene. Scale bar 10μm.

It is known that loss of repair/recombination factors lead to checkpoint bypass during meiosis [42, 43, 55]. Based on the results presented above, we were tempted to examine if SIRT1 was essential to induce such quality control mechanisms, specifically in response to exogenous damage. We saw a dose-dependent increase in the ratios of diplotene to pachytene cells in *Sirt1^Δmeio^* mice compared to *Sirt1^WT^* (Fig 7C). Our results suggest that upon induced damage, unlike in the wild type, loss of SIRT1 may lead to bypass of the pachytene/recombination checkpoints. Interestingly, compared to non-irradiated cells, irradiated diplotene cells showed patches of ©H2AX and the number of these patches was significantly higher in *Sirt1^Δmeio^* cells compared to *Sirt1^WT^* (Fig 7D and 7E). Together, these indicated persistence of damage even in diplotene and further corroborated our earlier findings on the role for SIRT1 in activating and/or coupling molecular factors involved in repair and recombination. Hence, we propose that in response to exogenous damage, SIRT1 is required for eliciting checkpoint mechanisms, which needs to be addressed in the future.

## Discussion

In this study, we report the importance of a NAD^+^-dependent deacetylase SIRT1 in regulating meiotic progression. Our findings reveal that SIRT1 is required to couple synapsis and meiotic DSB repair/recombination, and its absence leads to defective DSB repair and altered recombination frequency. Besides being one of the first reports to highlight the role of SIRT1 in meiotic progression, our study posits that de-/acetylation of molecular factors that govern these processes is necessary to elicit checkpoints under both basal and induced DNA damage conditions.

Loss-of-function mutants of SIRT1 in testis have indicated that it is indispensable for spermatogenesis [17, 19-22, 24]. Absence of SIRT1 has been shown to induce apoptosis and loss of meiotic populations, for example when *Sirt1* was knocked out using the pre-meiotic *Stra8-Cre* [17]. While, recent reports have provided some insights into its role in post-meiotic phases [17, 22], its importance in meiotic progression (specifically given its high expression in spermatocytes) remains to be unraveled. Our study, which has employed *Spo11-Cre* to knockout *Sirt1* only in spermatocytes, clearly shows that it plays a key role in meiotic progression. Given that the meiosis specific knockout of *Sirt1* does not lead to any loss of cells, it is likely that the phenotypes observed in the previous reports [17, 20, 24] reflect a cumulative effect of absence of SIRT1 in pre-meiotic and meiotic stages. Importantly, since lack of SIRT1 during meiosis did not lead to gross perturbations in meiotic populations, this model enabled us to unravel its role in regulating core meiotic processes viz. repair and recombination.

It is intuitive to expect that coupling of molecular processes to cellular progression through meiosis would possibly be regulated by post-translational modifications of both core and regulatory components. For example, phosphorylation is known to orchestrate meiotic progression, including by impinging upon DSB induction/repair efficiency and crossover frequency [56]. Even though acetylation is now regarded as a predominant modification on several proteins [57-59], if/how de-/acetylation of repair/recombination machinery affects meiotic progression is poorly addressed. In this context, our study clearly illustrates that absence of SIRT1 causes global hyper-acetylation of specifically non-histone proteins, and brings to the forefront the need to further investigate the interplay between protein de-/acetylation and meiosis in mammals. Our study also paves way for future efforts to investigate the possible links between metabolic inputs and meiosis given that SIRT1 is a NAD^+^-dependent deacetylase.

One of the key highlights of our study is loss of coupling between synapsis and repair/recombination in *Sirt1^Δmeio^* mice (Fig 6D). Given the tight interplay between synapsis and recombination, and the fact that loss of meiotic components also cause synapsis defects [39, 42], the current understanding of recombination-mediated control of progression through meiosis is limited [38, 39, 42, 43, 60]. Studies using individual or combination mutants involving *Mre11^ALTD/ALTD^*, *Nbs1^ΔB/ΔB^*, *Atm ^−/−^*; *Spo11^+/−^*, *Trip13^mod/mod^*, *p53^−/−^* and *Chk2^−/−^* have provided insights into regulation of both synapsis and recombination check points, which together control progression through meiosis [42, 43]. In this regard, our results not only establish SIRT1 as a key driver of meiotic progression but also as an upstream regulator of meiotic checkpoints.

Specifically, loss of SIRT1 led to delayed repair as indicated by high ©H2AX in early-mid pachytene, which was further corroborated by abnormal retention of RPA in late pachytene. Notably, the numbers of RPA foci at early pachytene were comparable between *Sirt1^WT^* and *Sirt1^Δmeio^*, indicating that the phenotype was unlikely to be caused by excess DSBs. Additionally, shorter SC-axis length at late pachytene in *Sirt1^Δmeio^* mice indicated delayed progression though meiosis [60]. Moreover, the pivotal role played by SIRT1 in exerting control over DSB repair was evident from the phenotype of *Sirt1^Δmeio^* mice exposed to irradiation induced exogenous damage. *Sirt1^Δmeio^* spermatocytes had exaggerated retention of ©H2AX when compared to the control, symptomatic of deficient repair. Intriguingly, however, at 3Gy and 6Gy of irradiation, loss of SIRT1 led to bypass of pachytene-to-diplotene checkpoint, albeit with persistent damage as indicated by increased number of diplotene ©H2AX patches in *Sirt1^Δmeio^* mice compared to *Sirt1^WT^*. Therefore, our results together uncover a dual role of SIRT1 in not only coupling synapsis to repair/recombination, but also in activation of checkpoints following exogenously induced damage (Fig. 6D).

It was also exciting to find that meiotic loss of SIRT1 led to a significant increase in crossover frequency as indicated by enhanced number of MLH1 foci. This is consistent with previous reports wherein increased DSBs and/or defective repair have been associated with altered recombination frequency [49, 50]. Together, these are significant findings since PTM based mechanisms that elicit recombination and repair checkpoints are less understood. In this regard, we propose that SIRT1-dependent deacetylation might be involved in setting a threshold for activation of either of these checkpoints, under both basal and exogenously induced damage conditions. In the future, it will be exciting to not only investigate the interplay between SIRT1 and pathways that induce these checkpoints but also in general to address the relevance of de-/acetylation-mediated control of meiotic progression.

Our efforts to gain preliminary insights into possible SIRT1-dependent molecular mechanisms during meiosis revealed components of the MRN complex (Fig 6B). Although, SIRT1-NBS1 interaction is known [28, 51], our results clearly illustrate that SIRT1 interacts with and affects acetylation of other components of MRN complex as well. In this context, we would like to highlight that *Sirt1^Δmeio^* mice phenocopy *Mre11* and *Nbs1* hypomorphic mutants [39], and together with the molecular data, it clearly suggests that SIRT1-MRN interplay is critical for meiotic progression. In the future, it will be interesting to investigate the role of de-/acetylation in controlling activity/localization of MRN complex during meiosis. It is also likely that SIRT1 could exert control over other key players such as ATM, p53, CHK2 and TRIP13 to mediate a tight coupling of synapsis, repair and recombination.

In summary, we have discovered a novel function of SIRT1 in meiosis. Our findings further highlight the importance of identifying mechanisms that affect or regulate core meiotic components. Specifically, given that mutation of some of these core-components lead to meiotic arrest, our results demonstrate that regulatory post-translational modifications, as brought about by SIRT1 in this case, are key determinants of meiotic outcome both in terms of governing quality of germs cells and recombination frequency.

## Materials and methods

### Ethics statement

The procedures and the project were approved and were in accordance with the Institute Animal Ethics Committee (IAEC) guidelines.

### Housing, AH1 and AH2

Mice housed in two different animal facilities at ACTREC-Mumbai (AH1) and IISER-Pune (AH2) were used in this study. While this was done due to shifting of our mice colony, it provided us the opportunity to score for the robustness of the phenotype when mice were reared in different housing conditions. The molecular/cellular phenotypes described in this manuscript were consistent between AH1 and AH2, and results specifically obtained from either of the facilities have been clearly indicated.

### Mutant mice

All animals were maintained on 12-hour light/dark cycle and given *ad-libitum* access to standard chow diet. Pups were weaned from mothers at 25 d.p.p. and group-housed later. *Sirt1-Exon4^lox/lox^* mice were obtained from Jackson laboratories (Jax-mice-ID 008041) and *Spo11-Cre* mice were a kind gift from Prof. Paula Cohen, Cornell University, Ithaca, USA. *Sirt1-Exon4^lox/lox^* strain was isogenized to C57BL/6N background for ten generations. Testis specific knockouts of *Sirt1* were generated using the strategy as shown in Fig 1A.

### Mice genotyping

For determining the genotype of the mice, tail clips or seminiferous tubules were digested and PCR was performed using KAPA HotStart Mouse Genotyping Kit (KAPA BIOSYSTEMS, Cat No. KK7352). The following primer pairs were used for determining genotype-Sirt1 genotyping: FP: 5’-GGTTGACTTAGGTCTTGTCTG-3’, RP: 5’-CGTCCCTTGTAATGTTTCCC-3’, Spo11-Cre genotyping: FP: 5’-TGGGCGGCATGGTGCAAGTT-3’, RP: 5’–CCGTGCTAACCAGCGTTTTC-3’, Post Cre excision Sirt1 Genotyping: FP: 5’-AGGCGGATTTCTGAGTTCGA-3’, RP: 5’-CGTCCCTTGTAATGTTTCCC-3’.

### Meiotic chromosome spreads

Meiotic chromosome spreads were prepared as described earlier [61]. Briefly, testes were harvested and collected in PBS, decapsulated and tubules detangled. Short segments of the tubules were placed in hypotonic lysis solution (30mM Tris pH 8.2, 50mM Sucrose, 17mM citrate trisodium dihydrate, 5mM EDTA, 0.5mM DTT and 0.1mM PMSF) for 90 minutes. Tubule segments were then transferred to 100mM sucrose solution, pH 8.2 and were finely diced/chopped using forceps. These were spread onto slides previously dipped in 1% paraformaldehyde with 0.15% Triton X-100 and were dried overnight in a humidified chamber at 37°C. Slides were then washed twice in 1X PBS and Photoflo™ and fresh spreads were used to score for chromosome synapsis to avoid breaks during freeze-thaw.

### Metaphase chromosome spreads

Testes were decapsulated in PBS and detangled using forceps. Single cell suspension of testicular cells was obtained by treating seminiferous tubules in 0.5 mg/ml Collagenase A (Roche, Catalog No: 23324223) for 45 minutes at room temperature. Debris was removed by passing this through two layers of gauze. Cells were washed twice with 2.2% sodium citrate (isotonic) buffer, and the final cell pellet was resuspended and incubated in 0.9% sodium citrate (hypotonic) solution at 37°C for 20 minutes. The cells were fixed in Carnoy’s Fixative (methanol:acetic acid :: 3:1) and were then dropped onto pre-warmed (60°C) slides and allowed to spread. DAPI was used to stain the DNA.

### Antibodies

The following primary antibodies were used as indicated in the figures: anti-SCP3 (Abcam, ab15093 and ab97672), anti-SCP1 (Abcam, ab15087), anti-©H2AX (CST, 9718), anti-RPA (Abcam, ab109394), anti-pRPA (Abcam, ab76420), anti-Mre11 (Merck, MABE 1153), anti-MLH1 (Santa Cruz, 550838), anti-TRF1 (Abcam, ab-1423-100), anti-SIRT1 (Merck, 07-131), anti-pan acetyl (CST, 9814S), anti-H3K9Ac (Diagenode, C15410004), anti-H4K16Ac (CST, E2B8W), anti-H3 (CST, CS-135-100) and anti-H4 (Abcam, ab7311). The following secondary antibodies for either immunofluorescence or immunoblot analyses: Goat anti-Rabbit IgG (H+L) Secondary Antibody, Alexa Fluor 488 (Invitrogen, A-11034), Goat anti-Mouse IgG (H+L) Secondary Antibody, Alexa Fluor 488 (Invitrogen, A-11001), Goat anti-Rabbit IgG (H+L) Secondary Antibody, Alexa Fluor 594 (Invitrogen, A-11012), Goat anti-Mouse IgG (H+L) Secondary Antibody, Alexa Fluor 594 (Invitrogen, A-11005), Anti Mouse IgG peroxidase antibody (Sigma, A4416), Anti Rabbit IgG peroxidase antibody (SIGMA, A0545) and Anti ArHm IgG peroxidase antibody (Abcam, ab5745)

### Immunofluorescence

Immunofluorescence was performed using previously described methods [42, 61]. Briefly, slides were washed in PBS + Photoflo™ and PBS + Triton X-100, blocked for 30 minutes and incubated overnight with indicated primary antibody at 4°C or with secondary antibodies for 45 minutes, and counter stained with DAPI. Spreads were washed after incubations with antibodies as described (36). The spreads were then imaged using Apotome epifluorescence microscope (Zeiss) and images analysed using Image J (Fiji). For quantification of RPA, pRPA and MLH1, only those foci that colocalized with SYCP3 axis were considered.

### Protein lysate preparation and immunoblot analyses

Protein lysates were prepared by homogenizing cells/tissues, and incubating on ice for 20 minutes, in either Radioimmunoprecipitation assay (RIPA) buffer (50mM Tris chloride, pH 8.0; 150mM Sodium chloride; 0.1% SDS; 0.5% sodium deoxycholate; 1% Triton X-100; 1mM sucrose) or TNN buffer (50mM Tris pH7.5, 150mM NaCl and 0.9% NP-40). Commercially available protease inhibitors PIC (Roche, Catalog No: 04693159001) and PMSF (Roche, Catalog No: 000000010837091001) and phosphatase inhibitors PhoStop (Roche, Catalog No. 04906837001) were added to buffer immediately before use. The cell debris was removed by centrifugation at 12,000xg for 15 minutes at 4°C and the protein concentration in the supernatant was determined by BCA assay. RIPA lysates were used for immunoblot analyses and TNN lysates were used for immunoprecipitation or immunoblot analyses. Immunoblots were developed using Chemiluminescence detection kit (ThermoFischer Scientific, Catalog No. P134080) and visualized using GE Amersham Imager 600. Band intensities were quantified using ImageJ.

### Histone Extraction from testis tubules

For extraction of histones from testis tubules, the protocol was followed as described earlier [62]. Briefly, tubules were homogenized in TNN buffer, followed by treatment of the pellet with 4N H_2_SO_4_ at 37°C. Histones were precipitated using Trichloroacetic acid (TCA), followed by washes with acetone. The pellet was finally resuspended in water and boiled in Laemmli buffer.

### Histological analyses of testis sections

Testis fixed in Bouin’s solution (Sigma, Catalog No. HT10132) were processed for obtaining paraffin embedded sections as per standard procedures. 5μm thick sections were used for staining. For Hematoxylin and eosin staining sections were deparaffinized and rehydrated before staining with Gill’s hematoxylin (Sigma, Catalog No. S076) and Eosin (Sigma, Catalog No. S007), as per standard procedures. Slides were finally treated with 95% ethanol, 100% ethanol and xylene and mounted in DPX.

### TUNEL assay

TUNEL assay for scoring apoptotic cells was performed using *In situ* cell death detection kit fluorescein (Roche, Catalog No.11684795910), as per manufacturer’s protocol.

### Gamma irradiation

3-month old mice (*Sirt1^Δmeio^* or *Sirt1^WT^*) were subjected to different non-lethal doses of Gamma Irradiation using Cobalt-60 source, as indicated. Immediately after the irradiation the mice were administered water with antibiotic (Meriquin-Enrofloxacin oral solution, 0.1% in autoclaved water) and were sacrificed after 72 hours for further analyses.

### Flow cytometry analysis

Single cell suspensions were obtained by Collagenase-A treatments as described earlier and were fixed in 70% ethanol at −20°C overnight. Following treatment with 300μg/ml RNase-A (Roche) they were stained with 25μg/ml propidium iodide (Sigma) and the populations of n, 2n, 4n were scored using FACS Fortessa (BD Biosciences) and analyzed using BD FACSDIVA 6.0 software.

### Cell culture and treatment with SIRT1 inhibitor

HEK293T cells were maintained in DMEM High glucose medium (Sigma, Cat No. D777) supplemented with 10% FBS and antibiotic-antimycotic, under standard conditions. They were treated with either EX527 (working conc. 10μM), inhibitor of SIRT1 or 0.1% DMSO for 16 hours. Cell pellets were used to obtain TNN lysates for immunoprecipitation or immunoblot analyses.

### Immunoprecipitation and interaction analyses

TNN lysates from cells/tissues were incubated overnight at 4°C with indicated antibodies and normal IgG was used as a control, and as described earlier [18]. Immune complexes were pulled down with Protein-G/Protein-A beads, as appropriate. For identifying SIRT1 interactors in testis endogenous SIRT1 was immunoprecipitated from six testes (three C57/BL6 mice). The complexes, eluted in 2X Laemmli buffer, were run on 12% SDS-PAGE through the stacking gel and the run was stopped once they reached the resolving gel. The gels were stained with Coomassie blue dye and the stained gel plugs were cut and washed with water/acetonitrile (50/50), reduced, alkylated and trypsin digested and processed for mass spectrometry analysis.

The extracted peptides were run on nanoLC-MS/MS with an UltiMate 3000 RSLCnano system (Dionex) coupled to an Orbitrap-Velos mass spectrometer (Thermo Scientific). Five microliters of each sample were loaded on a C-18 precolumn (300 μm inner diameter × 5 mm; Dionex) in a solvent made of 5 % acetonitrile and 0.05 % trifluoroacetic acid, at a flow rate of 20 μl/min. After 5 min of desalting, the precolumn was switched online with the analytical C-18 column (75 μm inner diameter × 50 cm; Reprosil C18) equilibrated in 95% solvent A (5 % acetonitrile, 0.2 % formic acid) and 5 % solvent B (80 % acetonitrile, 0.2 % formic acid). Peptides were eluted using a gradient of solvent B from 5 to 25 % in 80 min, then 25 to 50% in 30min, and 50 to 100% in 10min, at a flow rate of 300 nl/min. The mass spectrometer was operated in a data-dependent acquisition mode with Xcalibur software. Survey MS scans were acquired in the Orbitrap on the 350–1800 m/z range with the resolution set at 60,000 and AGC target at 1 × 106 ions, the 20 most intense ions were selected for CID (collision-induced dissociation), and MS/MS spectra were acquired in the linear trap with an AGC target at 5 × 103 ions, maximum fill time at 100 ms, and a dynamic exclusion of 60 s to prevent repetitive selection of the same peptide. Triplicate technical LC-MS measurements were performed for each sample. The mass spectrometry proteomics data have been deposited to the ProteomeXchange Consortium via the PRIDE [63] partner repository with the dataset identifier PXD014075.

Raw MS files were processed with MaxQuant software (version 1.5.2.8) for database search with the Andromeda search engine and quantitative analysis, as described earlier [64] Data were searched against Mus musculus entries in the Swissprot protein database (release UniProtKB/Swiss-Prot 2017-01). Protein quantification was performed using the LFQ intensity metrics from the MaxQuant “protein group.txt” output, to compare proteins identified in SIRT1-immunopurified and control samples. An average intensity value was calculated for each protein from the intensity values of the 3 MS technical replicate runs. Intensities were log2-transformed, and imputation of missing value with noise was performed in the Perseus software.

To assess interactors that belonged to particular cellular processes, proteins/peptides specifically enriched in SIRT1-immunoprecipitates were analyzed using GO analyses tool (https://david.ncifcrf.gov/) and STRING database (https://string-db.org/).

### Statistical test analysis

All analysis was performed using GraphPad Prism 6.0 or Microsoft Excel. Statistical significance was determined using Student’s t-test (between groups) or Mann-Whitney-U test (for MLH1 foci and SC length quantifications). For LC-MS/MS data, statistical analysis was performed in Perseus by applying a Student t-test between SIRT1 and control groups, and a global permutation-based FDR of 5% to detect proteins significantly enriched in SIRT1-immunopurified samples. P-values <0.05, <0.01 and <0.001 are indicated by *, ** and *** respectively for all experiments.

## Acknowledgements

We acknowledge the following funding sources: IFCPAR (Indo French Centre for the Promotion of Advanced Research) 4503-1, Swarnajayanti fellowship (DST Govt. of India. Grant number DST/SJF/LSA-02/2012-13) and TIFR-DAE (Govt. of India. Grant number 12P0122). We thank Paula Cohen, Cornell University, USA for sharing the Spo11-Cre mouse with us. We thank Gauri Shembekar for help with MLH1 and SC length quantifications. We acknowledge Investissement d’Avenir program ProFI (Proteomics French Infrastructure, ANR-10-INBS-08) for providing assistance with mass spectrometry. We thank NCCS-Pune, Dr. Boppana Ramanamurthy and Mr. Suresh Basutkar for help with gamma-irradiation. Animal house staffs at TIFR-Mumbai, National Facility for Gene Function in Health and Disease (NFGFHD) at IISER-Pune and ACTREC are acknowledged for help with animal experimentation and ACTREC histology department for help with testis sectioning. We also thank Dr. Kalidas Kohale, Dr. Shital Suryavanshi, Dr. Sachin Atole, Dr. Sagar Tarate, and Ms. Ritika Gupta for the help with animal experiments.

## Author contributions

Project conceptualization, funding acquisition and Supervision: UKS. Investigation and analyses: HK, SD, AF, AGP, UKS. Manuscript writing and data formatting: HK, SD, UKS.

## Supporting Information

### Legends to Supplementary figures

**S1 Fig. Characterization of *Sirt1^Δmeio^* mice.**

(A-B) Genomic DNA PCR from testis and tail clip of *Sirt1^WT^* and *Sirt1^Δmeio^* mice, (A) upper band (900 bp) corresponds to the WT and the lower band (450 bp) corresponds to the floxed-out *Sirt1*, (B) upper band (750 bp) corresponds to the floxed locus and the lower band (550 bp) corresponds to the WT locus. (C) Flow cytometry-based quantification of spermatogenic cell populations based on their DNA content, from animals housed at two animal houses (AH-1 and AH-2). N=3 per genotype, quantifications from 30,000 cells per animal. (D) Representative testis sections stained with H&E, shows no difference in testis morphology. (E) TUNEL assay on testis sections to score for apoptosis, showing marginal difference between *Sirt1^WT^* and *Sirt1^Δmeio^* mice. Arrows represent TUNEL +ve cells. Scale bar 100 μm. (F-G) Spermatocyte spreads stained for SYCP3 (red) and (F) H3K9me3 (green) to mark centromeres and (G) TRF1 (green) to mark telomeres. Representative images of cells in diplotene stage are shown. Scale bar 10 μm.

**S2 Fig. Synapsis is not altered at prophase I and metaphase I in *Sirt1^Δmeio^* mice.**

(A) Representative images of sspermatocytes in late pachytene, stained for SYCP3 (red) and SMC3β (green). (B) DAPI stained metaphase spreads from *Sirt1^Δmeio^* mice. Scale bar 10μm.

**S3 Fig. Abnormal retention of γH2AX in *Sirt1^Δmeio^* cells.**

(A) Immunoblot of ©H2AX from testis lysate of *Sirt1^WT^* and *Sirt1^Δmeio^* mice, showing higher levels in *Sirt1^Δmeio^* cells, N=3 per genotype and quantifications from triplicate samples. (B) Percent pachytene cells with abnormal retention of ©H2AX. N=4 per genotype, n=308 for *Sirt1^WT^* and n=384 for *Sirt1^Δmeio^* mice. (C-H) Percent pachytene cells at (C,F) early, (D,G) mid and (E,H) late stages, from animals housed at two animal houses, AH-1 and AH-2, respectively; classified as described in Fig 3C.

**S4 Fig. Persistent DSB repair intermediates in pachytene cells.**

Representative images of spermatocytes at late pachytene, stained for SYCP3 (red) and pRPA (green). Scale bar 10μm.

**S5 Fig. Acetylation status of histones and MRE11 is SIRT1-dependent.**

(A) Representative immunoblot for acetylation of acid extracted histones from *Sirt1^WT^*and *Sirt1^Δmeio^* mice. (B) Immunoblot for acetylation of MRE11 immunoprecipitated from control cells and cells treated with SIRT1 inhibitor EX527.

**S6 Fig. Sirt1*^Δmeio^* cells are hypersensitive to genotoxic stress.**

(A-C) Percent pachytene cells, subjected to 3Gy irradiation, at (A) early, (B) mid and (C) late stages, classified as described in Fig 3C. (D-F) Percent pachytene cells, subjected to 6Gy irradiation, at (A) early, (B) mid and (C) late stages, classified as described in Fig 3C. N=3 per genotype, n=294 for *Sirt1^WT^* and n=239 for *Sirt1^Δmeio^* for 3Gy irradiation and N=3 per genotype, n=180 for *Sirt1^WT^* and n=166 for *Sirt1^Δmeio^* for 6Gy irradiation.

**S1 Table. Interactome of SIRT1 in mammalian testes**

